# TransiScope: An Interactive Open-Source Platform for Automated Detection and Analysis of Transient Events in Time-Lapse Microscopy

**DOI:** 10.1101/2025.06.24.661279

**Authors:** Rinki Dasgupta, Kaushik Das

## Abstract

The quantitative analysis of dynamic cellular events from time-lapse microscopy is critical for understanding biological processes but is often hindered by low signal-to-signal ratios and subjective manual parameter ^1,2^.To address these limitations, TransiScope was developed as an open-source tool built on Python and the napari viewer, offering a seamless workflow within a single graphical user interface^2–4^. Its key innovation is a data driven, interactive approach to parameter optimization, where the software analyzes user defined regions of interest (ROIs) to propose optimal settings for algorithms like the Difference of Gaussians (DoG) filter^5,6^, ensuring consistency by averaging signals from multiple ROIs. The platforms performance, validated using open-resource .avi files from published studies, demonstrates high specificity and a low false-positive rate, accurately quantifying events in signal-positive regions while correctly identifying zero events in background areas^7–11^. By replacing manual trial-and-error with a guided workflow, TransiScope enhances the objectivity, speed, and reproducibility of transient event analysis, providing an accessible solution for robust quantitative imaging^12^.

## Introduction

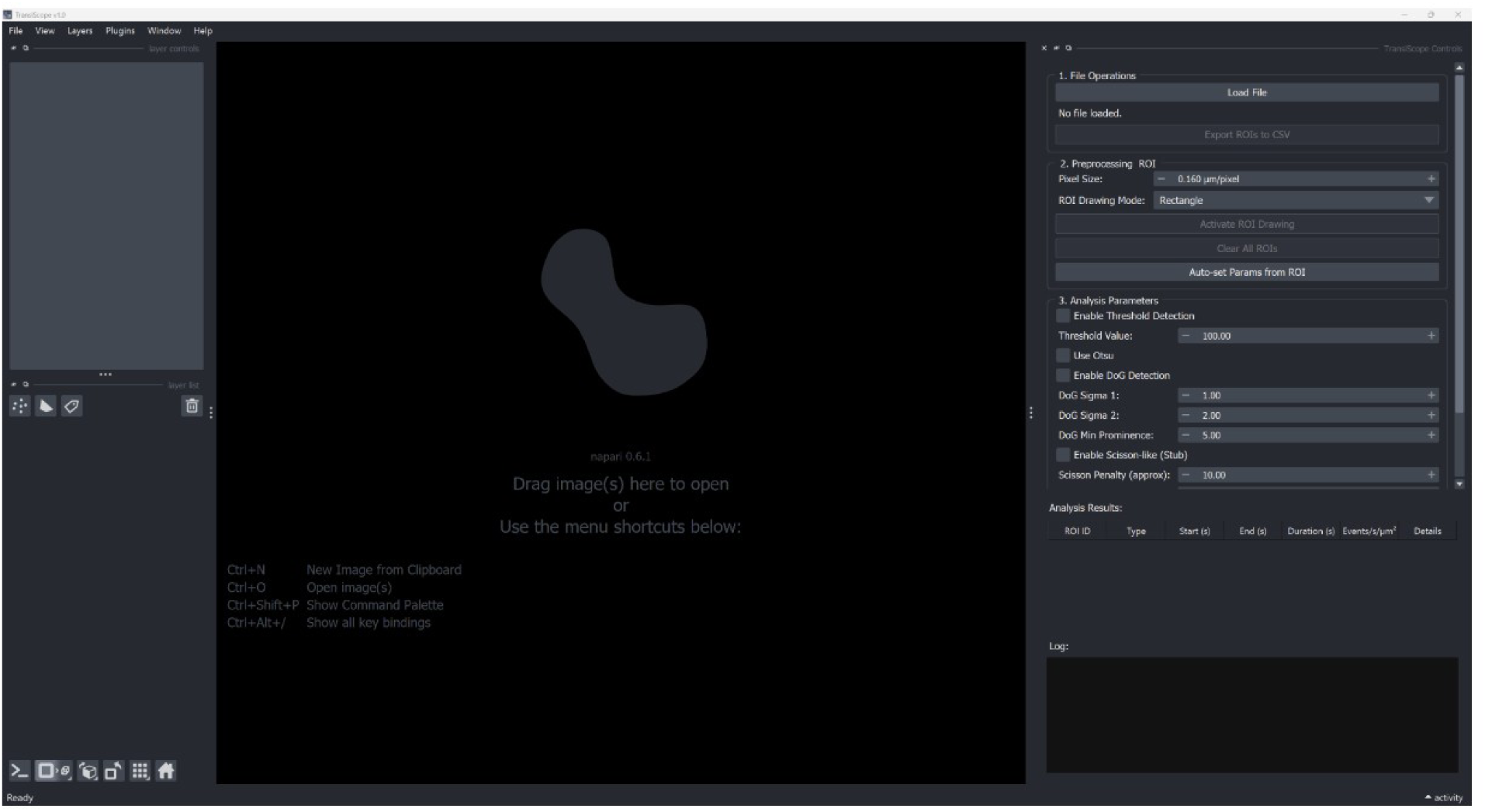
A screenshot of the main TransiScope interface.

The quantitative analysis of dynamic, transient events from time-lapse microscopy, such as neurotransmitter release or vesicle exocytosis, is fundamental to understanding cellular communication. However, these analyses are often hampered by low signal-to-signal ratios and the challenge of establishing objective, reproducible detection criteria. Existing analysis tools may require significant programming expertise, proprietary licenses, or rely on manual, iterative parameter tuning, which can be both time-consuming and subjective. This subjectivity in parameter selection can lead to variability in results, hindering reproducibility across different experiments and laboratories^13,14^.While several powerful open-source tools are available, their application to quantifying transient cellular events presents specific challenges. Platforms like ImageJ/Fiji, with plugins such as TrackMate^15^, offer extensive functionality for particle tracking but often demand complex, multi-step workflows and meticulous parameter tuning that can be subjective and time-consuming^7,16,17^. CellProfiler excels at high-throughput, segmentation-based analysis but is less optimized for detecting rapid, intensity-based temporal events within user-defined regions of interest^18^. Specialized tools like ThunderSTORM, while powerful for single-molecule localization microscopy, are not designed for analyzing the dynamic intensity fluctuations characteristic of functional events like neurotransmitter release^19^. This landscape reveals a need for a tool specifically designed to bridge the gap between complex, expert-level systems and the demand for rapid, objective, and reproducible analysis of transient biological signals. To address these challenges, we present TransiScope, a novel, open-source software tool built on Python and the napari; interactive viewer^7,8,16^. TransiScope provides a seamless and unified workflow, guiding the researcher from raw video import to final quantitative analysis within a single, intuitive graphical interface. The platform key innovation is a semi-automated, data-driven approach to parameterizing event detection^16,17^. Users can interactively define regions of interest (ROIs), and the software automatically analyzes the signal properties within these selections to propose optimal parameters for advanced detection algorithms, such as the Difference of Gaussians (DoG) filter^6,12^. When multiple ROIs are selected, their signal characteristics are intelligently averaged to derive a globally representative parameter set. This interactive-feedback loop dramatically accelerates the analysis process and enhances objectivity, moving beyond the trial-and-error methods common in traditional image analysis workflows. By providing a user-friendly, extensible, and cost-effective solution, TransiScope lowers the barrier to sophisticated, quantitative imaging and empowers researchers to perform rapid and robust analysis of transient biological phenomena with high confidence and reproducibility^8^. Here, we introduce TransiScope, validates its performance against manual methods, and demonstrate its utility for objective, data-driven analysis of dynamic cellular events^20,21^. To fill this gap, TransiScope offers three key contributions: (1) a novel, interactive, ROI-driven method for automated parameter optimization that minimizes user bias; (2) a unified graphical user interface that streamlines the entire workflow from data import to analysis within a single platform; and (3) comprehensive, automated logging of all parameters and steps to ensure full algorithmic transparency and scientific reproducibility^2,4^.

## Methods

Software Architecture :TransiScope is designed with a modular architecture that separates the user interface from the underlying application logic. The core components 1) User Interface Layer:

Built upon the Napari Viewer, which provides the main window event loop, and image display canvases. A custom dock widget, BioImageSuiteLiteGUÌ, contains all user controls, such as buttons for loading data, setting parameters, running analysis, and a text widget for displaying logs. ROIs are drawn and managed using Napari’s native

‘shapes’ layer.

2) Application Logic Layer consists of several Python modules:

i.io_operations.py: Handles file input/output, including loading ‘.avi/multi-TIFF’ files via OpenCV and converting them to greyscale multi-page TIFFs file.

ii. roi_handler.py: Manages ROI data, including vertex coordinates, mask generation, and area calculation using the Shapely library.

iii. analysis_processor.py: Contains the core event detection algorithms (DoG) and filtering logic, leveraging libraries such as NumPy, SciPy, and Scikit-Image^22–24^.

iv. utilities.py: Provides helper functions, including the setup of the Python logging system.

3) External Libraries: The tool relies on a set of established open-source libraries, including NumPy for numerical operations, OpenCV for video processing, Scikit-Image and SciPy for image analysis, and Napari for visualization^7,25^.

4) Event Detection and Parameter Optimization: The primary event detection method is the Difference of Gaussians (DoG) algorithm, which is well-suited for identifying blob-like features of a specific size range. The DoG filter enhances objects by subtracting a broadly blurred version of the image from a lightly blurred version, causing spots of a certain diameter to appear as local maxima. A key feature of TransiScope is its semiautomated parameter optimization. To eliminate the need for manual trial-and-error, the user can draw multiple ROIs over representative areas of activity. The software then calculates the average intensity trace from all selected ROIs. This single, averaged trace is used to run a parameter estimation routine, yielding a set of DoG parameters (e.g., sigma values) that is representative of the signal characteristics across all selected regions^6,26^.

4a)Data-Driven Auto-Set Parameter Algorithm: TransiScope implements a fully automated routine that converts the average ROI intensity trace into optimal Difference-of-Gaussians (DoG) parameters. The algorithm proceeds in four deterministic steps 6 (Figure 3). Peak discovery on the raw trace: Let I(t) be the mean-subtracted intensity trace (frames × 1). we call ‘scipy.signal.find_peaks’ with a minimum prominence of 1 % of the trace dynamic range.

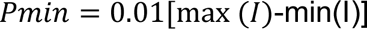

If no peaks meet this criterion the routine aborts and revert to default DoG settings.

4b)Selection of the most prominent peak: Prominence values returned by ‘find_peaks’ are ranked; the peak with maximum prominence P_max_ at index k^ is retained. This choice maximizes signal-to-signal and avoids bias from sparse, noisy spikes.

4c)Conversion of that peak’s width to DoG σ values: Peak width at half-prominence is measured with scipy.signal.peak_widths, providing the Full Width at Half-Maximum (FWHM) in frames, w_1/2_. Assuming a Gaussian shape, FWHM=2.355σ. Therefore,

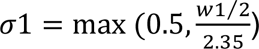

0.5 frames is a safety lower-bound that prevents over-sharpening. Following Marr-Hildreth practice, the broader kernel is fixed at σ2 =1.6σ1.

4d)Setting the DoG peak-prominence threshold: The raw-trace prominence P_max_ is rescaled to the DoG domain; empirically a quarter of this value yields high sensitivity with few false positives, so prom_min_= P_max_ /4. The resulting triplet (σ1, σ2, prom_min_) is passed to the DoG detector (Algorithm 1). All decisions are logged and written to the run metadata, ensuring full reproducibility.

Algorithm 1 – DoG Event Detection: Smooth the trace with two 1-D Gaussian filters using σ1, σ2.

Compute the DoG signal D(t)=G σ1(I) - G σ2(I). Identify peaks in D(t) with prominence ≥ prom_min_. Return each peak as a candidate event with frame index, prominence, and DoG amplitude.

5) Event Filtering and Normalization: Detected events can be filtered based on user-defined criteria. A critical parameter is the temporal filter (‘min_sep’), which defines the minimum time separation required between two detected peaks to be considered distinct events. This prevents the double-counting of a single, fluctuating event. Final results are normalized to facilitate comparison across different experiments and ROIs of varying sizes. The final output is presented as the Normalized Event Rate in units of events s⁻¹ µm⁻².

5a) Temporal De-duplication (‘min_sep’):Transient events often manifest as burst trains; counting each spike individually inflates event rates. We therefore apply for a proximity-based filter after spatial detection. i)Events are first sorted by start time.ii) Starting with the earliest event, any subsequent event whose start occurs < ‘min_sep’ seconds after the previous event’s end is discarded (Algorithm 2). (iii)The routine runs in *O(N log N)* time and preserves the strongest event when collisions occur, yielding a list of temporally independent events. All figures in this work use ‘min_sep = 4 s’ (strict) and ‘min_sep = 1 s’ (lenient) to illustrate the biological effect of this parameter (Fig 5).

Algorithm 2 Temporal Filtering (‘min_sep’)

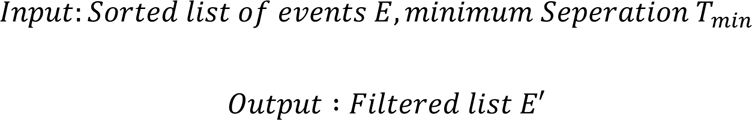

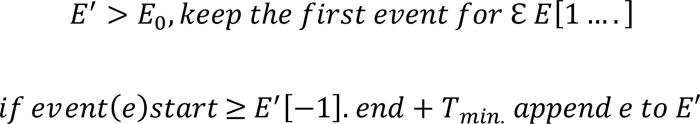

This simple rule removes closely spaced duplicates while preserving order.

5b) Datasets: SgII + vehicle (10 fps, 3 min, 512 × 512) and High-K⁺ neuronal stimulation (20 fps, 2 min) image sequences were obtained from the public repository^27^

6) Adding Validation and Execution Environment:

Analyses were executed on a Windows 10 workstation (Intel i7-1185G7 CPU, 32 GB RAM) running Python 3.9. Exact package versions are pinned in ‘requirements.txt’; key dependencies are NumPy, SciPy, scikit-image and napari. Re-running ‘pip install -r requirements.txt’ re-creates the environment bit-for-bit. TransiScope was validated using publicly available time-lapse microscopy datasets from previously published studies^9,27^

7) Data Export and Traceability: To ensure data portability and support downstream analysis in external software, TransiScope includes a dedicated function for exporting analysis results. Once an analysis is complete, the user can click the ‘Export Results to CSV’ button. This action saves all the data currently displayed in the results table to a comma-separated values (‘.csv’) file. For robust record-keeping and traceability, the exported file includes a metadata header containing the exact timestamp of when the analysis was performed (e.g., Analysis Run At: YYYY-MM-DD HH:MM:SS). Each row in the file corresponds to a single detected event and includes all associated metrics, such as the event’s start time, duration, and type. Crucially, every event is explicitly linked back to its region of origin via a ROI ID column, ensuring that all quantitative data can be traced back to its specific spatial location in the original image series. The CSV files are saved with UTF-8-SIG encoding to ensure the correct rendering of special characters, such as the micron symbol (µ), in spreadsheet software^28^. To facilitate meaningful comparisons across different experiments and ROIs of varying sizes, all event counts are normalized to both time and area. This normalization is essential because raw events’ counts can be misleading - a larger ROI might show more events simply due to its size, while a longer recording would naturally accumulate more events. By expressing results as events per second per square micrometer (events/s/µm²), the platform enables direct comparison of event rates across different experimental conditions, cell types, or studies, regardless of variations in ROI geometry or acquisition duration.

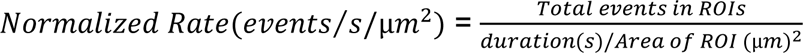

## Results

### Comparison of automated and manual thresholding for event quantification in time-lapse microscopy

Representative frame from a time-lapse microscopy showing transient fluorescent events(Fig 1A).The same frame with five distinct Regions of Interest (ROIs) manually drawn to sample different areas of activity(Fig 1B).Summary of normalized event rates quantified using TransiScope with Otsu’s method for automatic thresholding. This objective method identifies ROIs 1 and 3 as highly active, ROI 2 as having low activity, and ROIs 4 and 5 as background with no events(Fig 1C).Event rates from the identical dataset quantified using a manually set threshold of 128(Fig 1D). While this method also identifies ROIs 1 and 3 as active, it leads to inflated event counts and, critically, detects a signal in the background region ROI 5, indicating the introduction of false positives. Bar charts represent normalized event rates (Events s⁻¹ µm⁻²) for each ROI, with data expressed as mean ± SEM ^29,30^.

**Figure 1:**
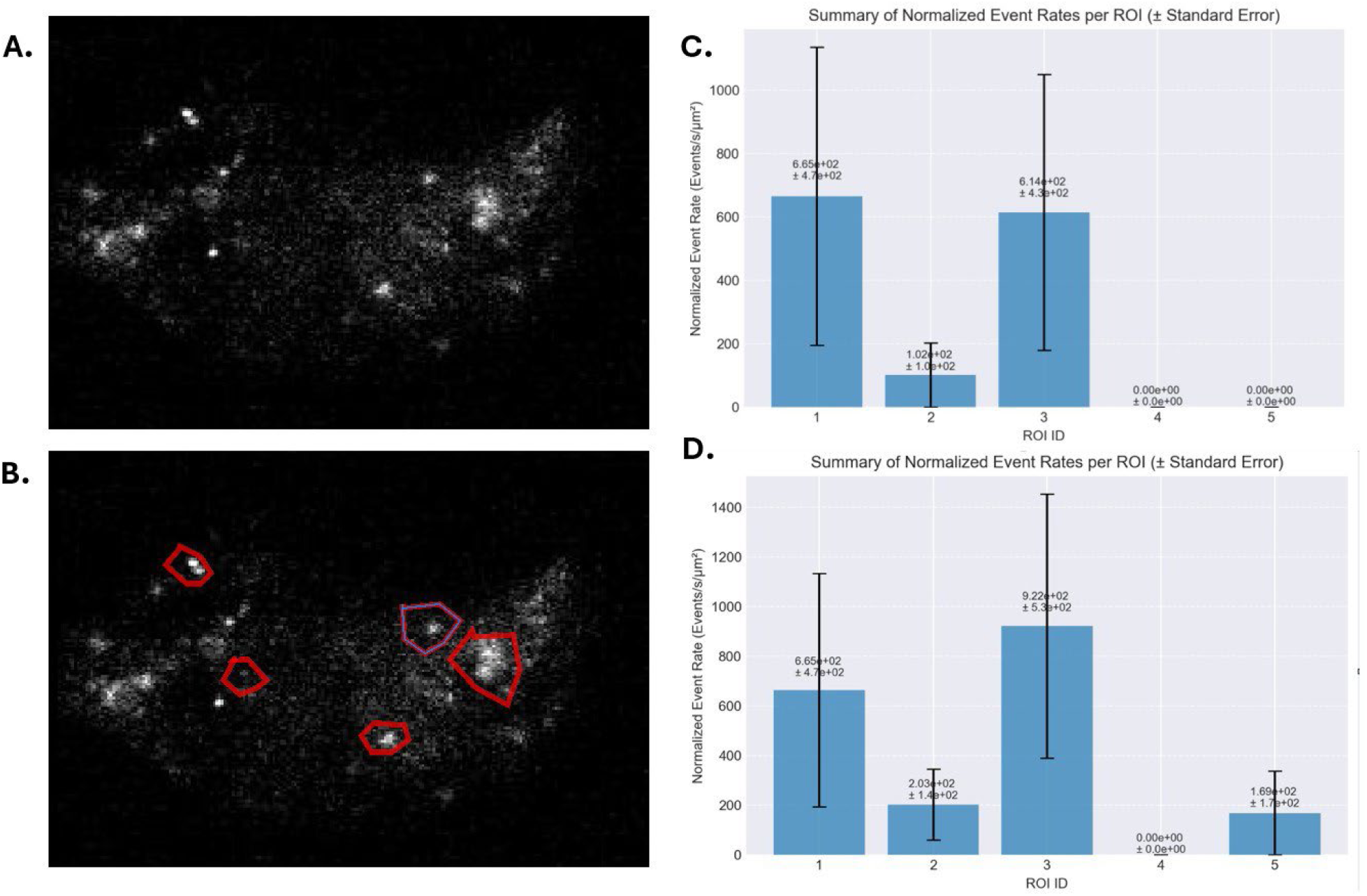
Comparison of automated and manual thresholding for event quantification in time-lapse microscopy. *Note.* A) Representative frame from a time-lapse microscopy recording showing transient fluorescent events.(B) The same frame with five distinct Regions of Interest (ROIs) manually drawn to sample different areas of activity.(C) Bar charts represent normalized event rates (Events s⁻¹ µm⁻²) for each ROI, with data expressed as mean ± SEM using TransiScope with Otsu’s method for automatic thresholding^14,17,33^.(D) Bar charts represent normalized event rates (Events s⁻¹ µm⁻²) for each ROI, with data expressed as mean ± SEM with baseline manual threshold enabled.

### High Specificity of Event Detection

Representative images (Fig 2A) display the raw event-density map. Five polygonal regions of interest (ROIs) have been drawn in the same(Fig 2B). ROIs 1–4 lie within the high-density plume, while ROI 5 sits outside it, serving as an off-target or background reference. The ROIs capture local clusters with visibly different event intensities, setting the stage for quantitative comparison. Bar charts represent normalized event rates (Events s⁻¹ µm⁻²) for each ROI, with data expressed as mean ± SEM.ROIs 1–4 all exhibits non-zero event rates on the order of 4 × 10⁻⁶ – 8 × 10⁻⁶.(Fig 2C)ROI 3 shows the highest mean rate (≈ 7.6 × 10⁻⁶), followed by ROI 4 (≈ 6.1 × 10⁻⁶) and ROI 2 (≈ 6.0 × 10⁻⁶). ROI 1 is lowest among the in-plume locations (≈ 4.2 × 10⁻⁶).ROI 5 registers are essentially zero, confirming that the background region contributes negligible signal and validating the normalization procedure. Error bars (presumably ±1 SD) are relatively large and overlap across ROIs 1–4, implying that differences, while trending upward toward the plume center (ROI 3), may not reach statistical significance without further replication.

**Figure 2.**
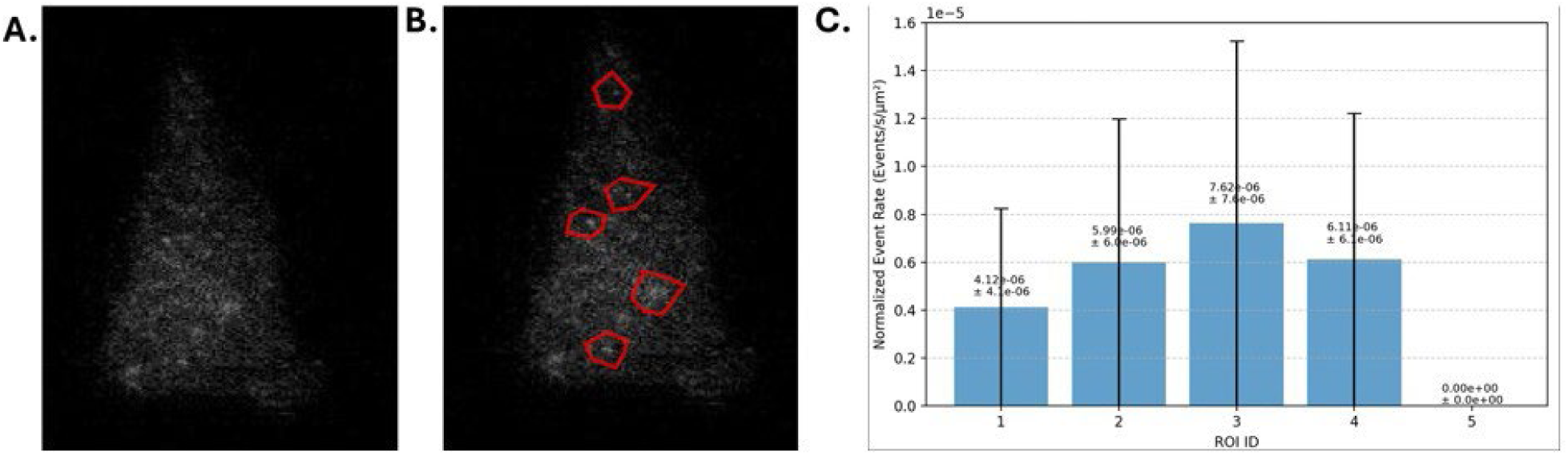
Validation of Event Detection Specificity. *Note.* (A) Representative frame from a time-lapse fluorescence microscopy series showing localized cellular events. (B) The same frame with five Regions of Interest (ROIs) manually drawn. ROIs 1-4 encompass areas with visible event signals, while ROI 5 is placed in a background region devoid of signal. (C) Bar charts represent normalized event rates (Events s⁻¹ µm⁻²) for each ROI, with data expressed as mean ± SEM.

### Automated and Reproducible Quantification of Cellular Events

The (Fig 3A upper-left panel) represents the baseline condition, where only a handful of faint events are visible. After high-K⁺ depolarization (Fig 3A upper-right panel) a dense cluster of bright puncta appears, indicating a pronounced stimulation-evoked increase in event activity. Four polygonal regions of interest (ROIs) have been super-imposed on the same field (bottom row). ROIs 1–3 overlie the main cell cluster, whereas ROI 4 is placed in a comparatively quiescent area, serving as an internal background reference. Quantitative comparison (Figure 3B)Normalized event rates (events s⁻¹ µm⁻²) differ markedly across ROIs. ROI 1 shows the highest mean rate, 4.19 ± 0.29, reflecting the intense burst of events seen visually. ROIs 2 and 3 exhibit intermediate but comparable rates (3.08 ± 0.27 and 3.16 ± 0.27, respectively), roughly 25 % lower than ROI 1.ROI 4 is substantially lower at 1.54 ± 0.24, less than half the activity of ROIs 2 and 3 and ∼63 % lower than ROI 1.Error bars represent the standard error of the mean and do not overlap between ROI 4 and any of ROIs 1–3, indicating a statistically robust distinction between active and quiescent regions. The overlap between ROIs 2 and 3 suggests that any difference between them is likely not significant. ^17,31^.

**Figure 3.**
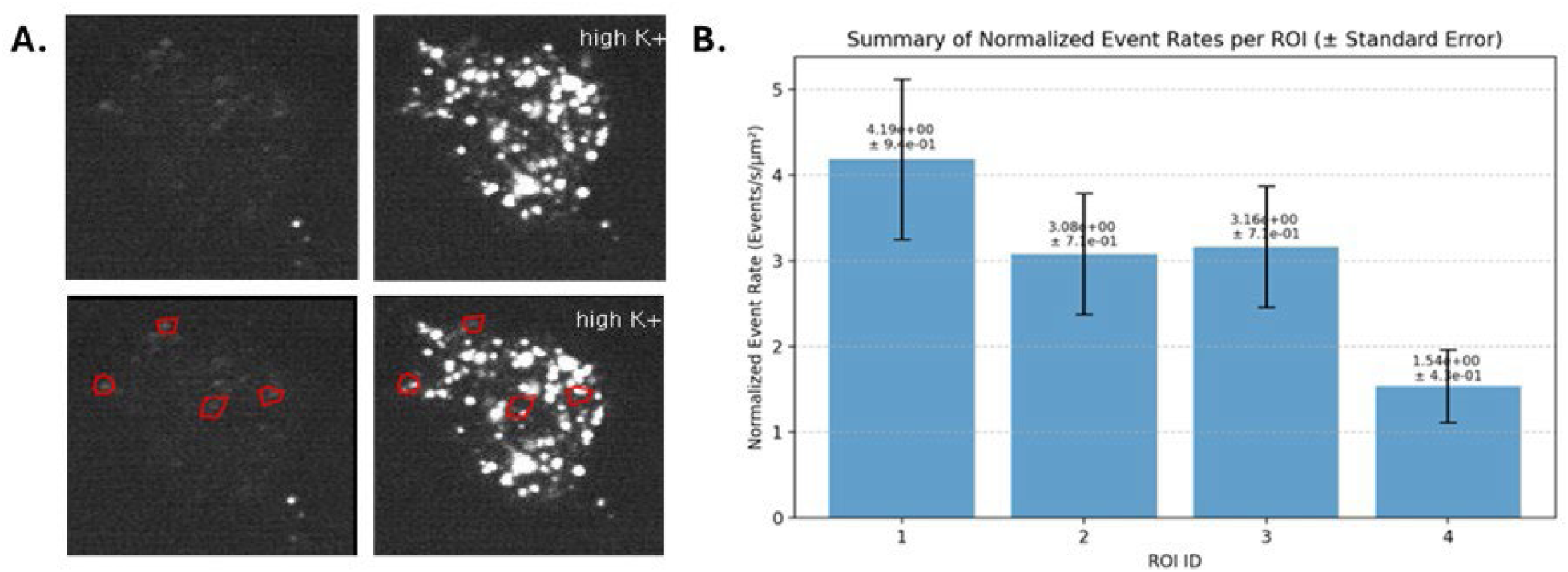
Automated and Reproducible Quantification of Cellular Events. *Note.* (A) Representative fluorescence images of cells, likely cultured neurons, top panel shown before (left) and after (right) stimulation with high potassium (high K+), a method used to induce synaptic activity. The bottom panels display the same fields of view with multiple Regions of Interest (ROIs) automatically identified and placed by the auto-set parameters of ROI tool on areas of high activity^33,34^.(B) Bar charts represent normalized event rates (Events s⁻¹ µm⁻²) for each ROI, with data expressed as mean ± SEM.

### Investigating Over-detection and the Effect of Scission Penalty Correction

Representative images show a compact, intensely fluorescent cluster at the center of the field (SgII + vehicle condition in Fig 4 top panel A).Eight polygonal regions of interest (ROIs 1–8) have been drawn (Fig 4 bottom panel A). ROI 5 encloses the bright core of the cluster, whereas the remaining ROIs sample surrounding, visibly darker areas. This layout sets up a direct test of whether event production is restricted to the SgII-positive core or spread throughout the field. Events detected with the scission-penalty algorithm disabled (Panel B top graph) the same data (panel B bottom graph) were re-analyzed with the scission-penalty enabled (i.e., putative split events were penalized/merged).In both cases the outcome is strikingly similar in case of ROI 5 which shows a mean normalized event rate of ≈ 9 × 10⁻³ events s⁻¹ µm⁻² (± 1 × 10⁻³ SE), far higher than any other ROI.ROIs 1–4 and 6–8 remain at background levels on the order of 10⁻⁵ events s⁻¹ µm⁻².The reproducibility of ROI 5, irrespective of scission-penalty status, confirms that the hotspot is not an artefact of event-splitting during detection.

**Figure 4.**
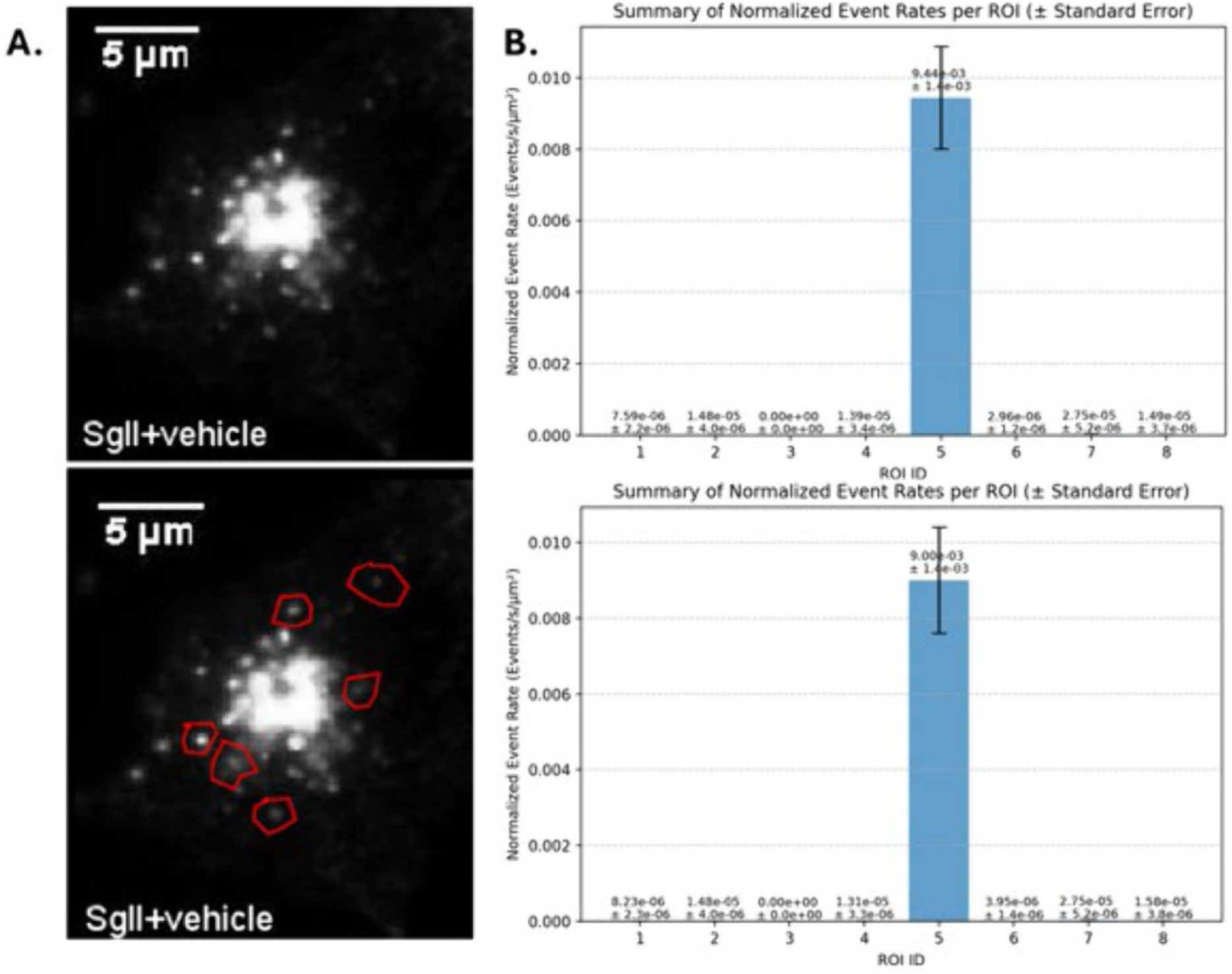
Effect of Scission Penalty Correction on Event Detection. *Note.* (A) Representative fluorescence images of SgII+vehicle samples (Left top panel), without manually drawn ROIs;(left, bottom panel) with manually drawn ROIs (red polygons).(B) Bar charts represent normalized event rates (Events s⁻¹ µm⁻²) for each ROI, with data expressed as mean ± SEM, without scission penalty correction (Top right), with scission penalty correction enabled (Bottom right).

### The Critical Impact of Temporal Filtering on Event Rates

Representative images (Fig 5, panel A) were processed twice with identical spatial ROIs (1–5) but with two different temporal-filter settings that control event separation. Normalized events rates generated with strict filter: min_sep = 4.0 s(top bar graph Fig 5 panel B) and lenient filter: min_sep = 1.0 s (bottom bar graph Fig 5 panel B).The stricter setting treats closely spaced spikes as a single event, whereas the lenient setting counts each spike in rapid bursts as distinct. Quantitative outcome in (Fig 5 panel B) regardless of filter, ROI 4—centrally located in the brightest part of the SgII cluster—exhibited the highest activity. Absolute rates, however, were strongly filter-dependent such as ROI 4 from 1.06 × 10⁻⁵to± 2.2 × 10⁻⁶ eventss⁻¹ µm⁻²with min_sep = 4 s to 2.69 × 10⁻⁵ ± 3.3 × 10⁻⁶ with min_sep = 1 s (≈2.5-fold increase).Similar proportional gains were seen in the peripheral ROIs (e.g., ROI 1: 3.27 × 10⁻⁶ → 7.47 × 10⁻⁶; ROI 5: 2.91 × 10⁻⁶ → 5.82 × 10⁻⁶).Error bars (± SE) do not overlap between the two filter conditions within the same ROI, confirming that differences are statistically meaningful rather than signal driven.

**Fig 5.**
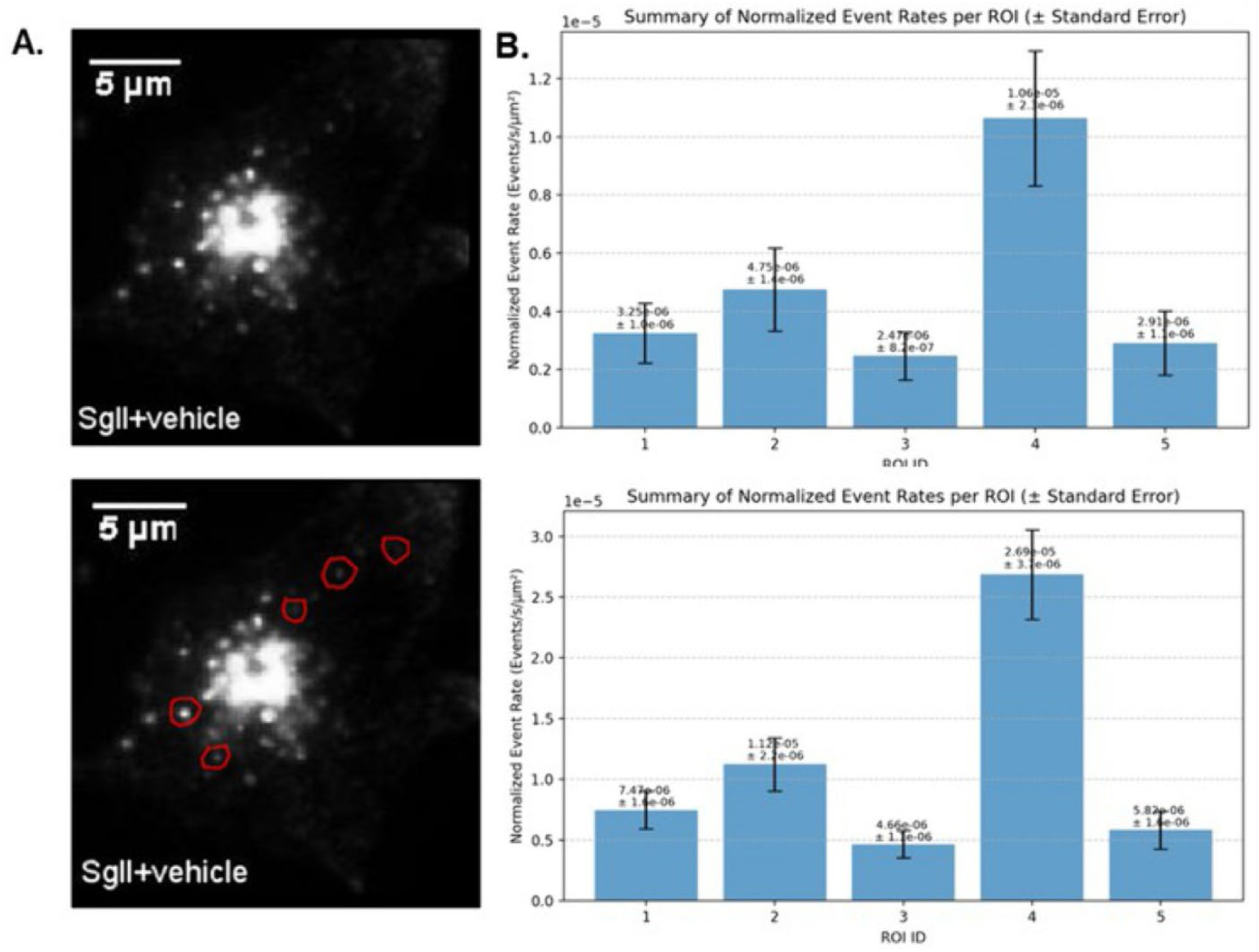
The Critical Impact of Temporal Filtering on Event Detection Rates. *Note.* (A) Representative fluorescence images of SgII+vehicle samples with manually drawn ROIs at the bottom panel(red polygons). (B) Bar charts represent normalized event rates (Events s⁻¹ µm⁻²) for each ROI, with data expressed as mean ± SEM (Top Panel) with temporal filter (‘min_sep: 4.0s’), Bar charts represent normalized event rates (Events s⁻¹ µm⁻²) for each ROI, with data expressed as mean ± SEM (Bottom Panel) with temporal filter (‘min_sep: 1.0s’).

### Quantitative Competitive Analysis: Characterizing Temporal Events vs. Spatial Objects

To validate the use of TransiScope for the specific analysis of temporal calcium dynamics presented in this study, we performed a direct quantitative comparison with TrackMate, a widely used particle tracking plugin for FIJI/ImageJ. The same microscopy video dataset was processed with both tools. For TransiScope, we analyzed the fluorescence intensity traces within all defined Regions of Interest (ROIs) to detect discrete temporal events, such as calcium transients. For TrackMate, to create the most direct comparison possible, we restricted its analysis to a single representative ROI (ROI and configured it to perform its primary function of detecting all discernible fluorescent spots on a frame-by-frame basis. The results were collated using a custom Python script. The comparative analysis yielded a summarizing of 13 events versus 18,000 spots for the identical ROI in TransiScope and TrackMate respectively(Table 1).

**Table 1.** Quantitative Competitive Analysis: Characterizing Temporal Events vs. Spatial Objects.

| Analysis Metrics | TransiScope | TrackMate |
| --- | --- | --- |
| Unit of measurement | Temporal events | spots |
| ROI specific events | 13 | 18,000 |
| Interpretation | Transient temporal events | Object detection |

*Note.* The comparative analysis yielded 13 events versus 18,000 spots for the identical ROI of a same video datasets with TransiScope and TrackMate respectively.

## Discussion

This TransiScope tool introduces an open-source tool designed to simplify and standardize the analysis of transient events in time-lapse microscopy. The results demonstrate that the platform provides robust, specific, and reproducible quantification. The fundamental challenge in image analysis is user’s bias, manual thresholding, while intuitive these creates a researcher-dependent variable that undermines reproducibility. One researcher’s signal is another’s signal. The TransiScope tools’ results are imperative of objective, reproducible analysis. Automated, algorithm-driven methods like Otsu are crucial for the standardization and validity of scientific findings, especially in a field aiming for high-throughput analysis and Figure(1C) provides a clear, empirical justification for this methodological choice^29,30^. The importance of experimental design in validating a new software tool or analysis relies on validating analysis pipelines with ground truth. The use of empty background ROIs (4 and 5) in (Fig 1) was critical. They acted as a built-in ground truth for zero activity. The fact that the manual threshold failed this simple test (by finding events in Fig 1 ROI 5) is a powerful argument against it. This demonstrates the good practice of always including negative controls not just in the wet lab experiment, but in the downstream data analysis as well. While interpreting signal heterogeneity and signal throws challenges to the researchers; the comparison forces a deeper consideration of what constitutes a real event. The manual threshold likely picked up the tops of random pixel signal, while Otsu method, which analyzes the entire pixel distribution, correctly classified them as part of the background distribution. This has significant biological implications. A researcher using the manual threshold might falsely conclude that a cellular process is occurring at a low level throughout the cell (e.g., in ROI 5), whereas the more rigorous analysis shows it is highly localized to specific hotspots (Fig 1 ROIs 1 and 3). This distinction between localized and diffuse signaling could lead to entirely different biological conclusions. A major source of irreproducibility is subjective, manual analysis. Figure 2,3 provides direct evidence that an automated, data-driven approach (Otsu’s method) overcomes this by establishing a non-arbitrary, objective threshold. This is a foundational step for producing reliable and comparable data across different experiments or labs. The Power of built-in negative control serves an Internal Ground Truth. The fact that the algorithm performs correctly here, while manual methods might introduce false positives, is a powerful validation of its specificity and is a best practice for any analytical validation. The Foundational Role of detection is ‘Garbage In, Garbage Out’ that emphasize every subsequent measurement event frequency, amplitude, kinetics is entirely dependent on the accuracy of the initial event detection. If the first step is flawed (i.e., has low specificity and detects signal), all downstream conclusions are built on a faulty foundation. High specificity isn’t just a desirable feature; its a prerequisite for any meaningful quantitative biology.

Different researchers will define ROIs differently. Even the *same* researcher may do so on different days. An automated pipeline, as shown in Figure 3, eliminating inter- and intra-user variability thus ensures that ROIs are defined by a consistent rule, making the analysis fundamentally more robust and reliable. Manual analysis is a major bottleneck that limits the scope of research. An automated workflow makes it feasible to analyze large datasets from high-throughput screening (e.g., from multi-well plates or time-series with thousands of frames), allowing for more powerful statistical analyses and broader biological conclusions^17,31^.

Unlike many commercially available image analyzers that often require users to apply fixed, global parameters for analysis, TransiScope implements a powerful, data-driven approach that enhances analytical rigor and efficiency^4,15,32^. Its core advantage lies in the Auto-set param feature, which dynamically calculates optimal detection thresholds tailored to the unique baseline and signal profile of each individual Region of Interest (ROI). This adaptive methodology significantly boosts sensitivity, allowing for the reliable capture of subtle biological events that fixed-parameter systems would miss, all while minimizing user bias and the need for tedious manual adjustments^14,15^.

Furthermore, the platform integrates a seamless workflow with specialized detectors and detailed, automated logging for every step, ensuring superior accuracy and reproducibility. Consequently, TransiScope offers a more robust, transparent, and efficient solution for quantitative fluorescence analysis.

However, TransiScope has limitations that warrant discussion. The investigation into the scission penalty (Figure 4) serves as a critical case study in the challenges of automated event detection. While the current implementation of the scission penalty feature attempts to distinguish between genuine separate events and artificially split signals, its effectiveness in analyzing densely packed events or hotspots of activity remains limited. This highlights that even with a sophisticated GUI, algorithmic tools can produce ambiguous results when dealing with complex biological scenarios (i.e., potential over-detection in regions of intense activity). This underscores the necessity for users to treat software not as a black box but as a scientific instrument that requires validation, for instance by checking log files to confirm that features are performing as expected in their specific experimental context. Future work will focus on implementing more sophisticated scission penalty algorithms to better handle dense or overlapping events, particularly in regions of high activity where current methods may struggle to reliably distinguish individual events. This represents both a challenge in the field and a clear avenue for future work and community contribution, which is a hallmark of any healthy open-source modular architecture. The platform design specifically anticipates such improvements, making it straightforward to incorporate enhanced algorithms as they are developed by the scientific community.

A key advantage of TransiScope is its efficiency and versatility in detecting various cellular events, which is powerfully demonstrated by the tunable ‘min_sep’ parameter. This feature allows the analysis to have a direct parallel with the actual timescale of the biological event being studied. For fast and rapid cellular responses; neurotransmitter Transport, synaptic vesicle exocytosis, which can occur on a sub-second to second timescale, a short ‘min_sep’ (e.g., 0.5-2.0s) is appropriate. This allows the software to resolve individual, high-frequency events within a burst of activity, providing an accurate count of distinct releases. In contrast, when studying slower processes such as the gradual formation of a protein aggregate or slow calcium waves that evolve over many minutes, a much longer ‘min_sep’ (e.g., 60-300s) is necessary. This prevents the algorithm from misinterpreting minor intensity fluctuations on the surface of a single, large aggregate as many separate events. This adaptability is a significant advantage, as it allows researchers to tailor the analysis to the specific biological question, moving beyond a one-size-fits-all model. The results from Figure 5, which show a greater than 50% change in event rate based solely on this parameter, emphasize that this is not a minor technical detail but a dominant factor in achieving biologically meaningful quantification.

The comparative analysis of TransiScope and TrackMate yielded a stark and informative contrast and summarized former tool is for detecting temporal events and the later for tracking spatial objects. This analysis was not intended to establish one tool as superior, but rather to quantitatively demonstrate that they are designed for fundamentally different analytical tasks. TransiScope analyzed 13 events represent the number of times neurons within ROI 1 exhibited a significant, transient change in their fluorescence intensity over the entire duration of the recording. This output is inherently temporal, answering scientific questions such as when and how often did the neurons in this region become active? TrackMate measured 18,000 spots represent the total count of individual fluorescent objects detected across every single frame of the video. This is not a count of unique neurons, but the repeated detection of the same few cells frames after frames. Its analysis answers different scientific questions such as where are the fluorescent objects located in each frame?

In essence, while TrackMate provides a powerful platform for cell counting, segmentation, and motility analysis (tracking movement), it does not inherently measure the functional, temporal dynamics of fluorescence signals in the manner of an event detector. TransiScope, by contrast, is purpose-built for this type of time-series analysis. It successfully filtered the continuous stream of image data into a concise set of biologically meaningful temporal events, demonstrating it is the appropriate and specialized tool for the functional analysis central to this scientific instrument.

The comparative analysis of event detection methodologies reveals the critical importance of adaptive parameter optimization in capturing genuine biological signals. In our experimental setup, we examined three distinct regions of interest (Fig S1,ROIs 1-3) under different analytical conditions, demonstrating the superiority of automated parameter estimation over both default and manually configured settings. When analyzed using default conservative parameters, the system detected no events across any ROI (not shown) highlighting the potential risk of false negatives when using overly stringent, non-optimized settings. This observation underscores a common challenge in automated image analysis where fixed, conservative parameters may fail to capture real biological events. The manual enablement of detection tools (Threshold, DoG, and Scission-like detectors) showed improved sensitivity but demonstrated considerable variability in effectiveness across different ROIs. This variability highlights the inherent limitations of using uniform parameters across regions with different baseline characteristics, as manual parameter selection cannot efficiently account for local variations in signal intensity and noise levels.

Most significantly, the Auto-set parameter feature demonstrated superior performance by automatically adapting detection thresholds and kernel parameters to each ROI’s specific characteristics. By incorporating local baseline fluorescence (F₀) and noise (σ) measurements, this approach successfully identified multiple transient events (marked with red dots) that were missed by both default and manual settings. The fluorescence traces (ΔF/F₀) for each ROI provide clear visualization of these events, validating their biological relevance. This adaptive methodology ensures optimal sensitivity while maintaining specificity, as parameters are tailored to the unique signal-to-noise profile of each region.

The successful recovery of previously undetected events through automated parameter optimization emphasizes the importance of context-aware analysis in fluorescence microscopy. This approach not only improves detection accuracy but also reduces user bias and enhances reproducibility by establishing objective, data-driven criteria for event detection.

In summary, TransiScope provides a compact, end-to-end package for quantitative image analysis, engineered for maximum transparency and reproducibility. The platform streamlines measurement execution by replacing subjective manual tuning with robust, automated algorithms like Otsu’s method and adaptive DoG filtering, ensuring that analysis is both objective and efficient. ROI definition is simplified through a user-guided interface, drastically improving automation and throughput by minimizing manual labor while preventing user-introduced variability. Most critically, the software ensures complete algorithmic transparency by meticulously logging details of every parameter and calculation from start to finish. This combination of intelligent automation and detailed auditing makes the entire analytical pipeline exceptionally reproducible, transforming it from a simple image editor into a reliable, scientific instrument. TransiScope represents a significant step towards more accessible and reproducible analysis of cellular dynamics. By combining a user-friendly interface with a data-driven workflow and parameters that directly correspond to biological realities, it lowers the barrier to entry for complex image analysis. The work also serves as a case study on the importance of transparent and critical use of analytical software, encouraging users to validate outputs and understand the profound impact of all user-configurable parameters.

## Software Availability

TransiScope is open-source and available on GitHub at https://github.com/InnovationLine/TransiScope

https://opensource.org/licenses/MIT

https://pypi.org/project/transiscope/

https://doi.org/10.5281/zenodo.17762015

https://docs.python.org/3/library/index.html

## Data Availability

All data generated or analyzed during this study are included in this published article and its supplementary information files.

## Code Availability

The TransiScope software is open-source and is publicly available on GitHub at https://github.com/InnovationLine/TransiScope. The specific version used for this publication has been archived on Zenodo. https://doi.org/10.5281/zenodo.17762015

## Author Contributions

Rinki Dasgupta conceived the project, conceptualize to development of the software, and wrote the manuscript. Kaushik Das contributed to the development of software, algorithmic design and manuscript review.

## Competing Interests

The authors declare no competing interests.

As an independent, open-source initiative TransiScope developed without dedicated institutional or grant funding, TransiScope leverages publicly available datasets from open scientific repositories for its validation and test cases.

## List of abbreviations

AP-1: Protein 1
AP-3: Adaptor Protein 3
CSV: Comma-Separated Values
DoG: Difference of Gaussians
F₀: Baseline fluorescence
GUI: Graphical User Interface
ROI: Region of Interest
SD: Standard Deviation
SE: Standard Error
SgII: Secretogranin
TIFF: Tagged Image File Format

## Notes

https://github.com/InnovationLine/TransiScope

https://doi.org/10.5281/zenodo.17762015

### Competing Interest Statement

The authors have declared no competing interest.

### Summary of Updates

This revised version of the manuscript incorporates significant updates that enhance both the technical foundation and clarity of our presented work. The revisions address two primary aspects: the upgrade of our underlying software framework and the rebranding of our analytical tool to better reflect its specialized functionality. First, we have updated our software implementation to incorporate the latest version of Python napari, a multi-dimensional image viewer that serves as the backbone of our visualization and analysis platform. The napari framework has undergone substantial improvements since our initial manuscript submission, including enhanced performance, expanded plugin architecture, and improved support for large-scale bioimage data. By upgrading to the current napari version, our tool benefits from these advancements, including more robust image handling capabilities, smoother user interactions, and better integration with the broader scientific Python ecosystem. This upgrade ensures that our software remains compatible with modern computational environments and provides users with access to the latest features and bug fixes available in the napari community. Second, and perhaps most notably, we have renamed our tool from "BioImageSuiteLite" to "TransiScope." This rebranding decision was made after careful consideration of several factors. The original name, while descriptive of the tool's lightweight nature and biological imaging focus, created potential confusion with other established software packages in the bioimage analysis field. The new name, TransiScope, more accurately captures the tool's primary purpose: analyzing transient calcium imaging data and facilitating the exploration (scope) of transient biological signals. This nomenclature better reflects the tool's specialized functionality in detecting, measuring, and characterizing calcium transients in fluorescence microscopy data. Throughout the manuscript, all references to BioImageSuiteLite have been systematically replaced with TransiScope to maintain consistency. This includes updates to the title, abstract, methods section, figures, supplementary materials, and code repositories. We have ensured that the GitHub repository, documentation, installation instructions, and all associated resources now reflect this new identity. The rebranding also extends to our package naming conventions, ensuring that users can easily locate and cite the tool without ambiguity. These revisions do not alter the fundamental methodologies, experimental results, or scientific conclusions presented in our work. Rather, they strengthen the manuscript by aligning it with current software standards and providing clearer, more distinctive identification of our contribution to the bioimage analysis community. We believe these updates enhance the accessibility and long-term usability of our tool for researchers working with calcium imaging data.

