## Supplementary materials for "TransiScope: An Interactive Open-Source Platform for Automated Detection and Analysis of Transient Events in Time-Lapse Microscopy"

**Fig S1***.* Adaptive parameter estimation is critical for robust detection of synaptic vesicle fusion events.

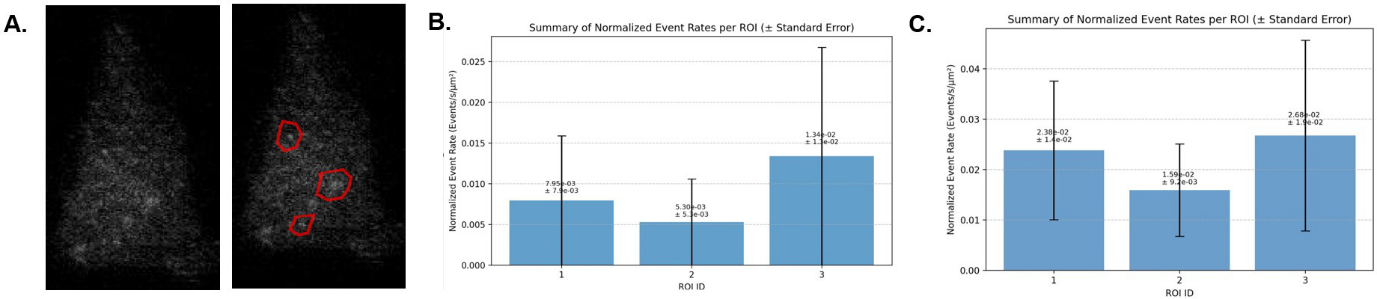

*Note.* (A) Representative maximum intensity projection of the time-lapse recording showing three manually selected Regions of Interest (ROIs 1–3).( B) Representative Bar charts shows normalized event rates (Events s⁻¹ µm⁻²) for each ROI, with data expressed as mean ± SEM manually enabled Threshold, Difference of Gaussians (DoG), and Scission-like detectors but without automated parameter estimation.(C)  Bar charts represent normalized event rates (Events s⁻¹ µm⁻²) for each ROI, with data expressed as mean ± SEM with Auto-set parameters enabled,.
